# Amoebic gill disease (AGD) in Atlantic salmon investigated through a holo-omic lens

**DOI:** 10.1101/2025.02.13.637958

**Authors:** Eiríkur Andri Thormar, Clémence Fraslin, Morten T. Limborg, Diego Robledo

## Abstract

Interactions between host genetics and the resident microbiota are complex. Understanding these interactions offers interesting alternatives for addressing gill health and disease resistance in salmonids. Amoebic gill disease (AGD), caused by *Neoparamoeba perurans,* remains a threat to Atlantic salmon, particularly in aquaculture settings where prevention and treatment options are scant. Selective breeding or genetic engineering towards increased AGD resilience present viable prevention strategies. While a number of studies have addressed AGD resistance in Atlantic salmon using transcriptomic and quantitative genetic approaches, the role of the Atlantic salmon gill microbiota in influencing AGD resilience and susceptibility needs further investigation. Addressing this, we leveraged a holo-omic approach using 16S rRNA bacterial profiling, and quantitative genetics, by considering the microbiota as an extended resistance trait of the salmon. We investigated the microbiota of AGD challenged Atlantic salmon in terms of two common resistance indicator traits: gill score and amoebic load. Subsequently we performed a GWAS using both the traditional indicator traits and traits of the microbiota. We found that the gill microbiota of the AGD affected salmon in this study was dominated by two bacterial families *Simkaniaceae* and *Arcobacteracea*. We found that microbial diversity and the relative abundance of *Simkaniaceae,* potentially derived from the amoeba, varied moderately with indicator traits. Additionally we identified several genomic regions that showed suggestive association with gill score and traits of the microbiota, and explored potential candidate genes for AGD resistance. Although much is still unclear regarding gill microbiota dynamics in gill disease, this study highlights the potential of addressing AGD through an integrative approach that considers the interplay between host genetics, the microbiota, and their roles in disease resistance.

## INTRODUCTION

Salmonids are one of the most commercially important aquatic food products accounting for 20% of the total value of exported aquatic products in 2022 (FAO, 2024). Diseases that occur in salmonid aquaculture are therefore a major threat to production, continued growth and sustainability of the industry. One important factor frequently mentioned in the context of salmonid aquaculture disease is gill health (Boerlage et al., 2020; Herrero et al., 2018; Mitchell & Rodger, 2011; Oldham et al., 2016). The gills are not only the primary organ of respiration in salmonids, but also serve other important functions such as Ph regulation, excretion of nitrogenous waste, hormone regulation and ionic regulation (Evans et al., 2005; Koppang et al., 2015). Being exposed to the surrounding water, the gills present one of the first lines of defence against invading pathogens and thereby also a potential point of entry for infectious agents (Cabillon & Lazado, 2019; Koppang et al., 2015; Salinas, 2015).

Amoebic gill disease (AGD) is one of the major threats to gill health of marine salmonids and hence of major importance in salmon aquaculture (Oldham et al., 2016). Outbreaks of AGD can lead to severe economic losses and high mortality rates (Oldham et al., 2016) and in extreme scenarios mortality rates up to 80% have been recorded in Atlantic salmon sea cages (Steinum et al., 2008). The primary causative agent of AGD is the amoeboid *Neoparamoeba perurans* (Crosbie et al., 2012). AGD pathology presents itself by white lesions, increased mucus production, epithelial hyperplasia and respiratory disruptions (Adams & Nowak, 2003; Marcos-López & Rodger, 2020; Zilberg & Munday, 2000). Moreover, AGD is often accompanied by other infectious agents resulting in a more complex disease phenotype often termed complex gill disease (CGD) (Boerlage et al., 2020; Herrero et al., 2018). AGD prevention and treatment options are scant, and currently limited to freshwater or hydrogen peroxide bathing (Marcos-López & Rodger, 2020; Rodger, n.d.). Therefore other options such as selective breeding or genetic engineering to increase resilience of Atlantic salmon stock to AGD may present alternative prevention strategies. Understanding natural disease resistance mechanisms to AGD in Atlantic salmon will be an immensely valuable resource for taking steps towards prevention of AGD.

A number of studies have addressed AGD resistance in Atlantic salmon using transcriptomic and quantitative genetic approaches. (Robledo et al., 2018) identified two regions on chromosome ssa18 which indicated suggestive association with AGD resistance, (Aslam et al., 2020) identified SNPs on chromosomes ssa01, ssa02 and ssa05 which were significantly associated with AGD resistance and (Boison et al., 2019) identified regions on chromosome ssa04, ssa09 and ssa13. Literature suggests that AGD resistance has a polygenic nature, with heritabilities estimated in the range of 0.09-0.48–typically characterised as moderately heritable–where the estimates differ based on source population of Atlantic salmon and the number of re-infection cycles (Aslam et al., 2020; Boison et al., 2019; Kube et al., 2012; Lillehammer et al., 2019; Robledo et al., 2018; Taylor et al., 2007, 2009). Studies in which transcriptomic analyses have been conducted have revealed differential expression of genes involved in cell adhesion and proliferation, mucin secretion, cell cycle control, red blood cells, and immune response shedding further light on mechanisms of resistance (Boison et al., 2019; Botwright et al., 2021; Marcos-López et al., 2018; Robledo et al., 2020; Wynne et al., 2008). Another layer may be added to the context of AGD resistance – that of the gill microbiota.

The microbiota is an often overlooked layer in the context of the host and has been exploited relatively little in the context of disease resistance. Recently (Schaal et al., 2022) showed a potential link between host genetics, metabolic rate and gut microbiota in AGD, demonstrating that including the layer of the microbiota is a potentially valuable asset in terms of disease resistance. Several studies have addressed the microbial communities of Atlantic salmon gills in the context of AGD (Birlanga et al., 2022; Botwright et al., 2021; Bowman & Nowak, 2004; Clinton et al., 2024; Downes et al., 2018; Slinger et al., 2020, 2021). It appears that the gill microbiota is generally quite dynamic–not unexpected given its close proximity to the surrounding water–but altered during an AGD outbreak, whether it be the presence or altered relative abundance of known pathogenic taxa or differences in diversity. It has further been suggested that some bacteria (*Winogradskyella* sp. in this case) may be able to enhance the severity of an AGD infection (Embar-Gopinath et al., 2005, 2006). Moreover bacteria have been reported to live in close association or as endosymbionts of the amoeba itself in cultures (Benedicenti et al., 2019; MacPhail et al., 2021; A. Nylund et al., 2018), and even differences in microbial composition among virulent and attenuated amoeba cultures has been described (Ní Dhufaigh, Botwright, et al., 2021; Ní Dhufaigh, Dillon, et al., 2021). Thus these complex dynamics of the host, parasite and microbiota are challenging to address, especially in context of host genetics and disease resistance.

Looking at this complex system in a more holistic way–namely the host and the associated microbiota in the same framework–may have potential in shedding further light on mechanisms of AGD resistance. Holo-omics which integrates combined -omics layers from host and its associated microbiota (Nyholm et al., 2020; Odriozola et al., 2024) provides a unique framework for investigating host-microbiota interactions in the holobiont–the host and its microbiome together (Baedke et al., 2020; Bordenstein & Theis, 2015; Zilber-Rosenberg & Rosenberg, 2008)–while offering new insights into ecology and evolution (Alberdi et al., 2022; Foster et al., 2017; Nyholm et al., 2020; Theis et al., 2016) and has great potential to be applied in industries such as aquaculture (Limborg et al., 2018).

We intended to investigate whether we could leverage a holo-omic framework to shed more light on AGD resistance/susceptibility. Here we utilise a combined approach of 16S metabarcoding and quantitative genetics to address AGD resistance. We characterise the gill microbiota of Atlantic salmon subjected to an AGD disease challenge, and investigate microbiota composition in relation to traditional AGD phenotype scoring methods (gill score and amoebic load). We then performed a genome wide association study (GWAS) to investigate SNPs associated with the phenotypic scoring methods and the microbiota.

## METHODS AND MATERIALS

The AGD challenge data used in this study has been published and described Robledo et al., 2020. Briefly, a cohabitation AGD challenge was performed using 797 Atlantic post-smolt salmon (∼18 month, mean weight after challenge 463.5g) from 132 nuclear families from a commercial breeding programme (Landcatch, UK). For the cohabitation challenge, fish were infected from an ongoing in vivo culture and fish with a similar level of AGD infection (measured as gill damage) were used as seeder in the cohabitation challenge with a ratio of 15% seeder to naive fish. The experimental challenge was conducted at the experimental facilities of University of Stirling’s Marine Environmental Research Laboratory, Machrihanish (Scotland, UK) in a 4m3 seawater tank (water temperature between 13 and 14°C, at 33-35 ppt of salinity). The challenge consisted of three separate cycles of infection with a recovery period after two infections (Taylor et al., 2009). A fresh water treatment was applied 21 days after the start of the challenge followed by a week of recovery and by the addition of seeder fish. During the challenge the fish were checked visually four times daily and in the third cycle of infection the disease was allowed to progress until sampling point.

At sampling fish were terminated by an overdose of anaesthetic (Phenoxyethanol, 0.5mg/L) and gill damage was recorded for both left and right gills by a single operator. Gill damage was scored from 0 for clear gills (healthy red gills, no gross sign of infection) to 5 for heavy AGD infection level (Extensive lesions covering most of the gill surface) according to (Taylor et al., 2016), the mean score of the two gills was used as resistance phenotype. One gill was dissected out and stored in ethanol for amoebic load estimation by qPCR using *N. perurans* specific primers and future 16S bacterial profiling.

### V3-V4 16S rRNA bacterial profiling

Out of 797 a total of 77 samples were used for 16S bacterial profiling, Equal sizes of gill tissue were cut off each sampled salmon and transferred to a lysing E-matrix tube containing 0.9ml of DNA/RNA shield (Zymo research). Prior to DNA extraction samples were lysed for 5 min in Tissuelyzer (Qiagen) at 30GHz after which they were spun down at 16.000g. In a randomised order, 400 µL lysate from samples was transferred to a 96 deep-well plate. DNA extraction was then performed using the Quick-DNA MagBead Plus kit (Zymo research) using the recommendations of the manufacturer. Negative extraction controls containing 400 µLDNA/RNA were included in the process. The DNA concentration was measured using a Qubit fluoromenter (Invitrogen). Prior to amplification a qPCR was performed on a subset of the samples to check for inhibitor presence and optimal cycle number. Initial amplification was performed using the forward and reverse primers 341F (5′-CCTAYGGGRBGCASCAG-3′) and 806R (5′-GGACTACNNGGGTATCTAAT-3′) (Yu et al., 2005) targeting the V3-V4 region of the bacterial 16SrRNA. Each amplification reaction consisted of 7.5 µL ddH2O, 9.5 µL AccuPrime SuperMix II (Invitrogen), 1 µL 10 µM forward 16S primer, 1 µL 10 µM 16S reverse 16S primer, and 1 µL sample DNA, totalling 20 µL. The amplification PCR settings were as follows: incubation at 95 °C for 10min, followed by 30 cycles of denaturation at 95 °C for 15s, annealing at 53 °C for 20s and extension at 68°C for 40s, after the completed cycles this was followed by an 10min final extension at 68°C. The Nextera XT index kit v2 Set A (Illumina) was used to uniquely index the PCR products. The indexed PCR products were then visualised on a 1% agarose gel and subsequently pooled in an equimolar fashion based on band strength and purified using SPRI beads. Libraries were quantified using the Agilent BioAnalyzer 2.100. Paired end sequencing was performed using the Illumnia MiSeq platform, reagent kit v3 at 600 cycles by the GeoGenetics Sequencing Core, University of Copenhagen, Globe institute. Negative extraction controls and PCR controls were included in the library construction and sequenced for downstream quality control.

### Bioinformatic processing and analysis of 16S amplicon sequencing data

Approximately 12.9 million read pairs were generated for the 83 samples and controls. Initial quality checks were performed using *FastQC* (Andrews, 2010) and *MultiQC* (Ewels et al., 2016). Adapter and initial quality trimming was subsequently performed using TrimGalore (Krueger et al., 2021). The *DADA2* pipeline (Callahan et al., 2016) was used for further quality filtering, trimming, and ASVs inference and removal of chimeric sequences. Taxonomy was assigned in the *DADA2* framework using *Silva/v138* database training set (Quast et al., 2013). *LULU* (Frøslev et al., 2017) was then applied for post clustering curation of the ASVs to minimise errors after which *Decontam* (Davis et al., 2018) was applied to remove contaminant ASVs from the dataset (Table S3 lists the contaminant taxa). The ASVs were further processed using the R-package *Phyloseq (McMurdie & Holmes, 2013)*. ASVs assigned to Eukaryota, Mitochondria or Chloroplast were removed and ASVs were then agglomerated on the family level based on lack of resolution on the genus level and to minimise noise. ASVs were normalised using simple proportions. A Hill numbers framework using the R-package *hilldiv* (Alberdi & Gilbert, 2019) was used to carry out alpha diversity analyses. Differential abundance analysis was performed using the Wilcoxon-rank sum test with an FDR adjusted p-value. Beta-diversity estimations were performed using a PCoA with Bray-Curtis distance and statistical differences estimated using PERMANOVA with Bray-Curtis distance from the R package *Vegan* (Oksanen et al., 2020). Statistical analyses were performed in *R/4.3.1* (R Core Team, 2023). Additional packages used for visualisation were *ggplot2* (Wickham, 2016), *ggpubr* (Kassambara, 2020), and *patchwork* (Pedersen, 2024).

### Genotyping and estimation of genetic parameters

The fish were genotyped using a SNP array with 47K SNPs (Houston et al., 2014), after DNA extraction from fin clip tissue samples using DNeasy 96 tissue DNA extraction kit (Qiagen, UK) as described in Robledo et al., 2018. Standard quality control was performed using *PLINK v1.90b6.21* (Purcell et al., 2007) where individuals with less than 95% of their SNPs genotyped were removed, SNPs with a minor allele frequency (MAF) less than 0.05, call rate of less than 95% and deviated from the Hardy-Weinberg equilibrium (p<10^-6^) were removed.

This resulted in 26,459 SNPs for 58 individuals. Next the software GCTA (Yang et al., 2011) was used to construct a genomic relationship matrix (GRM) with:

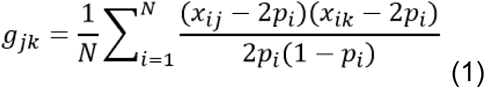

Where *g*_*jk*_ denotes the relationship between individual j and k. N refers to the total number of SNPs (26,459), the copy number of reference allele for a i^th^ SNP for the j^th^ and k^th^ fish is denoted by *x*_*ij*_ and *x*_*ik*_and *p*_*i*_ is the reference allele frequency.

The Average Information Restricted Maximum Likelihood (AI-REML) (Gilmour et al., 1995) algorithm applied in GCTA was then used for estimation of genetic parameters based on the GRM using the following linear mixed model approach:

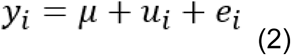

Where *y*_*i*_ denotes a phenotype for the i^th^ fish, μ is the mean in the population, *e*_*i*_is the random additive genetic value for individual i, following a normal distribution 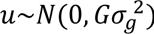, where G is the GRM made with GCTA (Eq.1) and 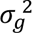 the estimated genetic variance. *e_i_* is the residual effect following a normal independent distribution 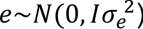 where 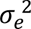 denotes the residual variance.

A GWAS was then performed in GCTA using a mixed linear model association with the leave-one-chromosome-out (loco) approach.

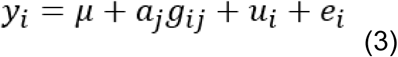

Where *y*_*i*_ denotes a phenotype for the i^th^ fish, μ is the mean in the population,for the j^th^ SNP *a*_*j*_ denotes the additive genetic effect of the reference allele with the genotype for individual i(*g*_*ij*_) represented with 0,1 or 2. *e*_*i*_is the residual effect following a normal independent distribution 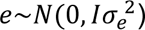 where 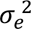 denotes the residual variance. *e* is then a random vector of polygenic effects following a normal distribution 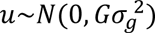, where G (because of the mlma-loco approach) is a partial GRM constructed with 28 chromosomes after leaving out the chromosome containing the j^th^ SNP, and 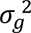 is the estimated genetic variance. A Bonferroni correction (α=0.05) was used to determine the genome wide significance threshold (−*log*10(α/*n*)) where *n* is the number of SNPs, and the chromosome wide significance threshold (−*logg*10(α/(*n*/29))). The phenotypes used were both the traditional AGD scoring methods (amoebic load and gill score) and traits of the microbiota which were; the unweighted and weighted alpha diversity measures, the first two axis of a beta diversity PCoA using bray curtis distance, and inverse normally transformed (INT) relative abundances of families that contributed over 3% of the total relative abundance in the dataset.

## RESULTS

### Sequencing and sample overview

A 16S amplicon library was sequenced consisting of DNA extracted from 77 gill samples including negative extraction and PCR controls and a positive control totalling 12.9 million read-pairs related to 83 samples. After quality filtering and trimming of sequences, ASV inference, taxonomic identification, removal of contaminant sequences and post clustering curation we were left with a dataset comprised of 844 ASVs. 35% of the ASV were not assigned on the genus level hence the ASVs were merged on the family level and analysed accordingly. After filtering out samples with a low ASV count and samples not assigned both Ct value for amoebic load and a gill score we were left with 59 samples containing a total of 180 bacterial families. Most of the samples had an assigned gill score around 3-4 while amoebic load Ct values were more evenly distributed among the 59 samples. For analysis of microbiota composition we grouped the samples by gill score into light-moderate (gill score <3.5) and moderate-advanced (gill score ⋝3.5) by degree of gill damage as described in (Taylor et al., 2016) and by amoebic load into Ct<30, 30<Ct<35, Ct>35 to appropriately cover the range of amoebic load Ct values.

### Microbiota profiles vary by AGD scoring methods

We first investigated the general composition of the microbiota on the family level, since 35.5% of the total abundance of ASVs was unassigned on the genus level. After filtering, we found that 61.44% of the overall relative abundance is attributable to two families namely, *Simkaniaceae* (31.3%) and *Arcobacteraceae* (29,79%). After *Simkaniaceae* and *Arcobacteraceae* the families *Vibrionaceae*, *Marinomonadaceae*, *Pseudoalteromonadaceae*, *Rhodobacteraceae*, *Flavobacteriaceae* each contributed over 3% of the overall relative abundance. The gill microbiota of the AGD challenged salmon in this study can therefore be described as dominated by two Families namely *Simkaniaceae* and *Arcobacteraceae* and a few families contributing between 1-3% of relative abundance (Figure 1G).

**Figure 1:**
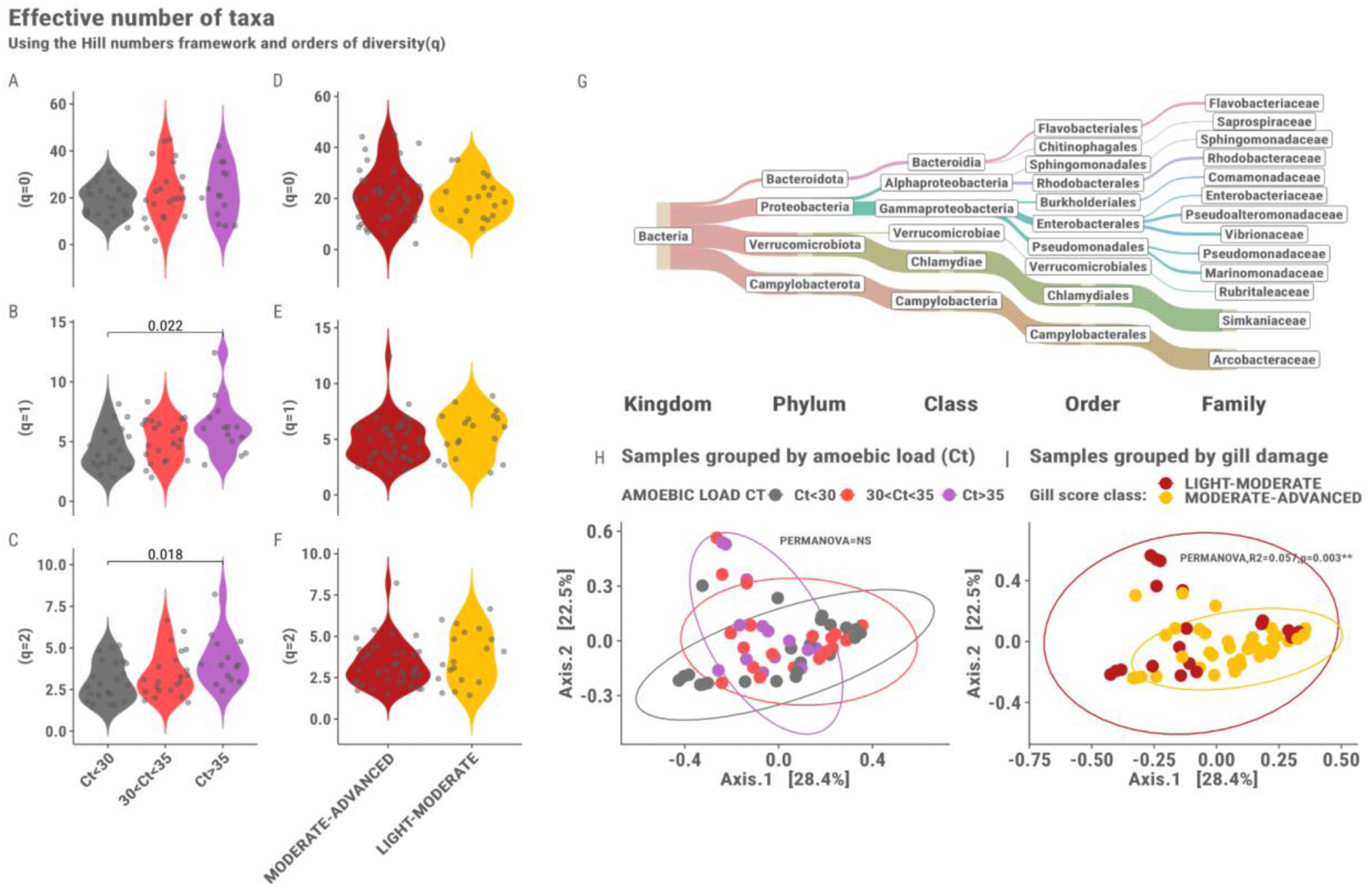
The left panel shows the comparison of alpha diversity estimates based on hill numbers order of diversity q=0, q=1 and q=2 among samples grouped by A-C) amoebic load Ct value where the colours represent Ct<30 (grey), 30<CT>35 (pink) and Ct>35 (purple), and D-F) gill damage where the colours represent the visually scored gill damage groups, Light-Moderate (yellow) and moderate-advanced (red). The top right panel G) shows a Sankey diagram of the bacterial families constituting over 1% of the overall relative abundance in the dataset where the thickness of the lines represents the relative abundance. The bottom right panels H-I) Show beta diversity PCoAs using Bray-Curtis distance of samples grouped by amoebic load Ct value and gill damage respectively.

We then looked at the diversity among the samples grouped by amoebic load and gill score. Using alpha diversity metrics with hill numbers and orders of diversity (Alberdi & Gilbert, 2019) we found significant differences in diversity between samples grouped by amoebic load on the Family level (Figure 1A-C). These significant differences were observed for the two orders of diversity q=1 (p-adjusted=0.024) and q=2 (p-adjusted=0.017). No significant difference was found on the ASV level (Figure S1). When we grouped samples according to severity of gill damage no significant difference was seen in alpha diversity on the family level (Figure 1D-F) or ASV level (Figure S1). Overall the weighted microbial diversity measures (q=1 and q=2) trend to increase with a lower amoebic load (Figure 1A-C). Moreover, The alpha diversity measures indicate that the samples were largely dominated by a few bacterial families confirming previous observations.

The beta diversity did not differ among samples grouped by amoebic load (Figure 1H-I) but differed slightly among samples grouped by gill damage (PERMANOVA,R2=0.057,p= 0.003). Considering the little difference in ordination space and a significant result in a test for homogeneity of dispersion(p=0.006), the effect observed may be due to differences in group dispersions.

### A Simkaniaceae bacterium correlates with disease status

Clustering of taxa abundances across samples revealed no general patterns in terms of amoebic load or gill damage (Figure 2A), but microbiota specific correlations. Differential abundance analysis of the bacterial families constituting more than 1% of the overall relative abundance shows that one bacterial family, *Simkaniaceae,* is differentially abundant (p.adjust=0.037) among samples grouped by gill damage (Figure 2C). No differentially abundant bacterial families were detected in samples grouped by amoebic load. Notably the representative ASV of the *Simkaniaceae* has a 99.78% percentage identity (using BLAST) with *Candidatus* Syngnamydia Salmonis, that may be an intracellular bacterium of the amoeba as it has been found in co-culture with the amoeba and identified in AGD before (A. Nylund et al., 2018; S. Nylund et al., 2015). Interestingly there is a significant correlation (R2=0.29,p=0.024) between the amoebic load Ct value and the relative abundance of *Simkaniaceae* (Figure 2B) suggesting a co-occurrence pattern with this bacterium and the amoeba.

**Figure 2:**
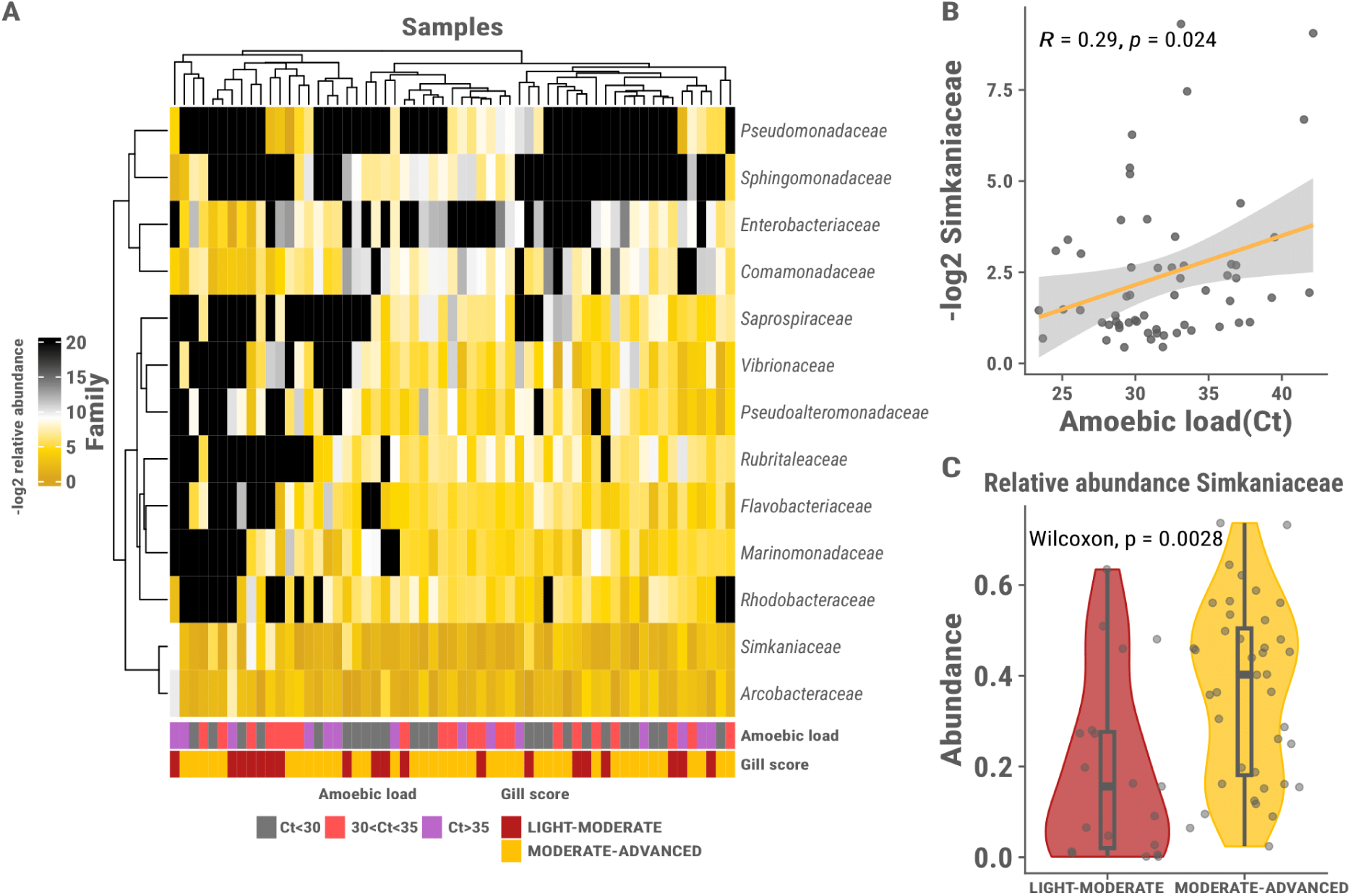
A) Shows -log2 transformed relative abundances of the bacterial families constituting over 1% of the total relative abundance across samples on a gradient from black (low) to dark yellow (high). The dendrograms represent the hierarchical clustering of samples and taxa abundances using euclidean distance and complete linkage. The bottom legends show the samples based on groupings of amoebic load Ct value and gill damage respectively. B) Shows the correlation between the -log2 transformed relative abundance of Simkaniaceae and amoebic load Ct value. C) Shows the differential abundance of Simkaniaceae in samples grouped by gill damage. The colours represent the gill damage groups, Light-Moderate (red) and moderate-advanced (yellow).

### GWAS identifies suggestive association with gill score and bacterial traits

We then performed a GWAS using both the traditional AGD scoring methods (amoebic load and gill score) as a phenotype together with chosen phenotypic metrics from the microbiota (see methods). One sample was discarded after filtering leaving 58 samples for the GWAS.

No meaningful inference could be made from heritability estimates due to large standard errors, likely due to the modest sample size (Table 1 for the phenotypes with suggestive SNPs, and Table S1 for all phenotypes tested). This is also reflected in the expected/observed p-value q-q plots where there is a slight deflation towards the tail of the q-q plots (Figure 3B). While no genome-wide or study-wide significant SNPs were found for any of the traits tested, we did detect some outlier SNPs at the suggestive significance threshold (Table 2).

**Table 1:** Variance component estimates for AGD resistance for phenotypes where suggestive SNPs were observed.

| Phenotype | $\sigma_a^2(\pm se)$ | $\sigma_p^2(\pm se)$ | $\sigma_e^2(\pm se)$ | $h^2(\pm se)$ |
| --- | --- | --- | --- | --- |
| **Gill score** | $0.07 \pm 0.1$ | $0.32 \pm 0.06$ | $0.26 \pm 0.11$ | $0.2 \pm 0.32$ |
| **Beta diversity axis.2** | $0.01 \pm 0.01$ | $0.04 \pm 0.01$ | $0.03 \pm 0.01$ | $0.17 \pm 0.35$ |
| **Alpha diversity(q=0)** | $13.21 \pm 31.44$ | $90.99 \pm 17.12$ | $77.78 \pm 33.09$ | $0.15 \pm 0.34$ |

$\sigma_a^2(\pm se)$ : The genetic variance, $\sigma_p^2(\pm se)$ : The phenotypic variance, $\sigma_e^2(\pm se)$ the residual variance, $h^2(\pm se)$ : the heritability estimate

**Figure 3:**
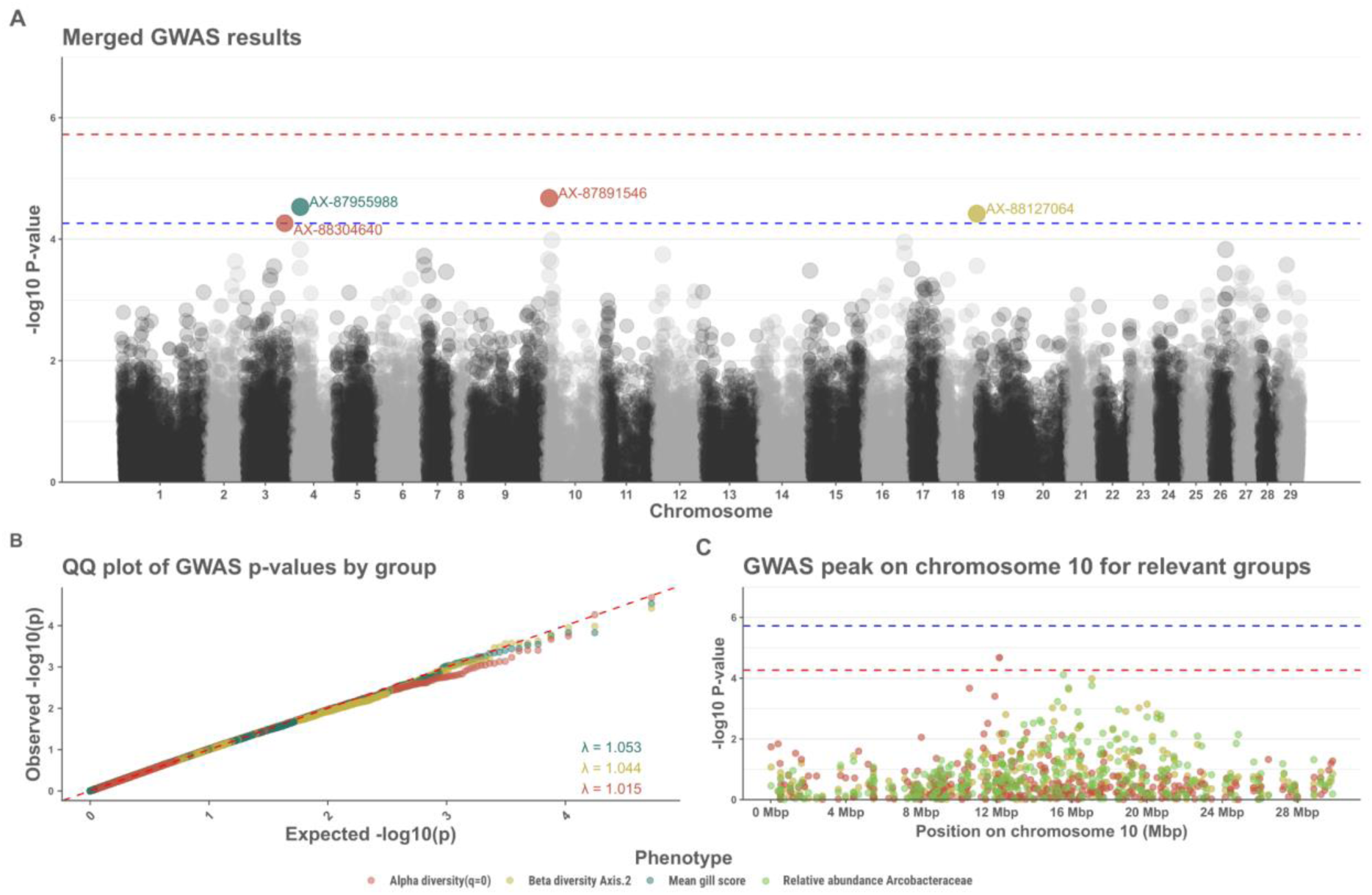
A) A merged GWAS for AGD resistance for the phenotypes with suggestive SNPs. The coloured dots represent the SNPs crossing the suggestive threshold and the colors represent the phenotype. The horizontal red line represents the genome wide significance threshold and the blue line represent the suggestive threshold B) q-q plot of the observed/expected p-values for the phenotypes with suggestive SNPs, the colors represent the phenotypes. C) A GWAS plot for the peak observed on chromosome 10, the colors represent the different phenotypes where the peak was observed.

**Table 2:**
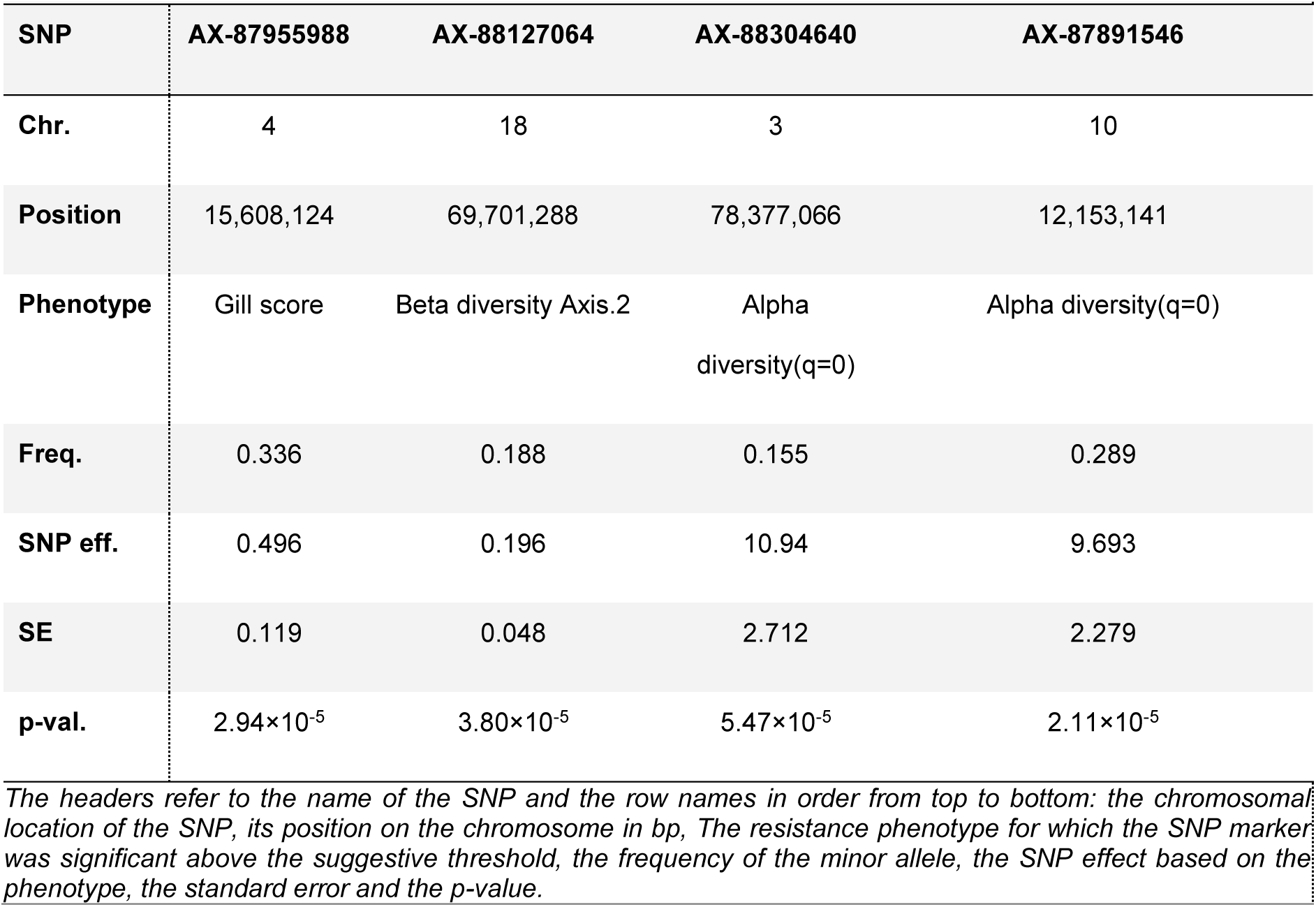
SNP markers reported to be significant at the suggestive threshold in the GWAS.

A single suggestive SNP (AX-87955988) was found on chromosome 4 when gill score was used as a phenotype (Figure 3A). Looking at the genotypes of that suggestive SNP we can see that individuals with the AA genotype tend to have a lower gill score compared to the individuals carrying the B allele (Figure 4A). Applying the microbiota diversity measures as phenotypes revealed a suggestive SNP (AX-88127064) on chromosome 18 for the 2nd axis of the beta diversity PCoA (Figure 3A). No obvious differences were observed on the 2nd axis on the PCoA other than individuals with the AA genotype seem to cluster more towards the lower end of the 2nd axis compared with individuals with the B allele (Figure 4B).

**Figure 4:**
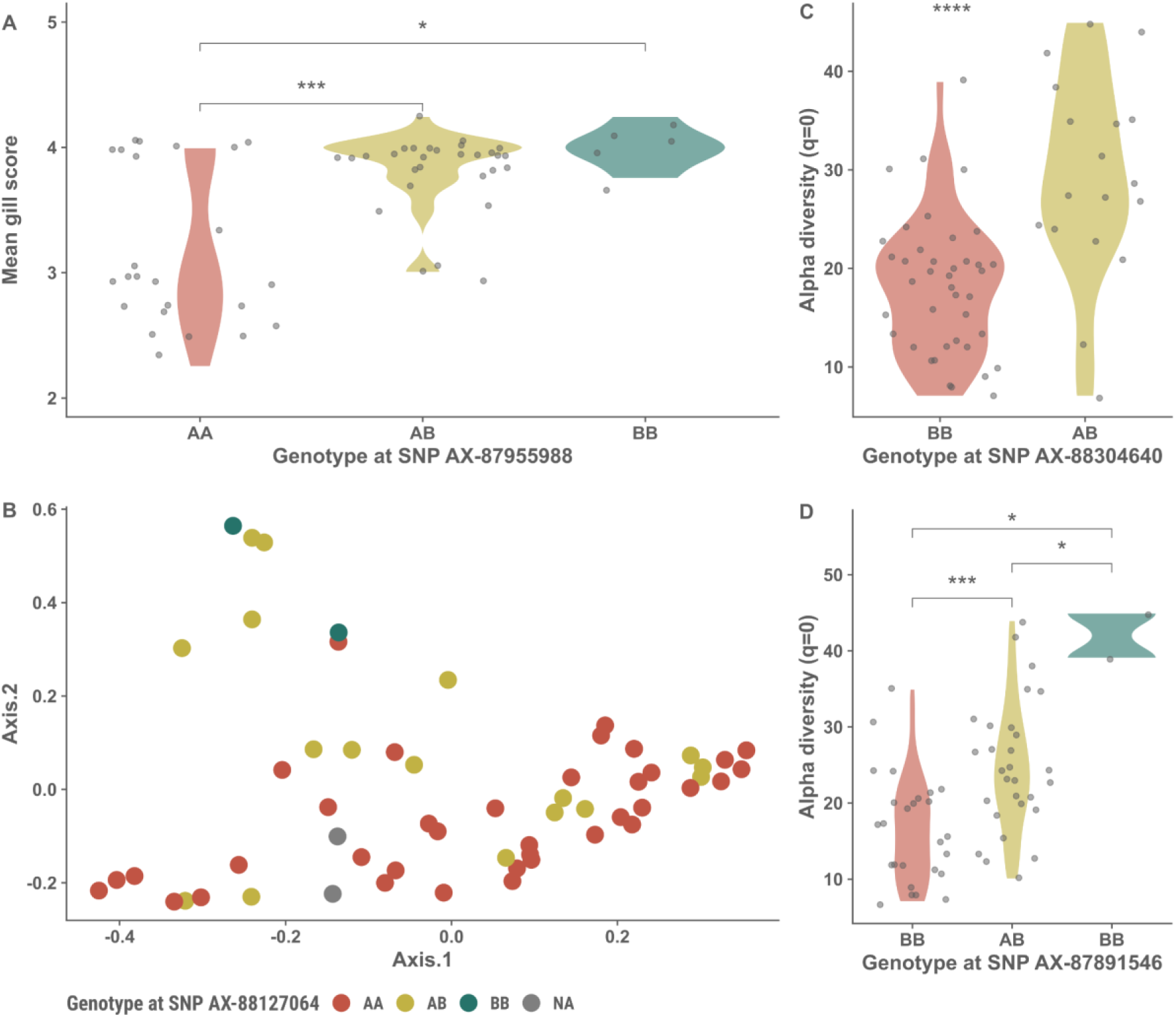
Violin plots and a PCoA depicting the differences in phenotypes among genotypes on the suggestive SNPS for A) Gill score on SNP AX-87955988, B) 2nd axis of a beta diversity PCoA on SNP AX-88127064, C)-D) Alpha diversity (q=0) on SNPs AX-88304640 and AX-87891546 respectively.

Two SNPs (AX-88304640 and AX-87891546) on chromosome 3 and 10 were significant at the suggestive threshold for richness estimates (Figure 3A). In both instances individuals with the BB genotype have lower diversity (at q=0) than do individuals carrying the A allele (Figure 4C-D).

No other suggestive SNPs were found for the relative abundances of the other bacterial families tested. Interestingly, a peak is present on chromosome 10 for both the 2nd axis of the PCoA and the richness estimates (Figure 3C). Furthermore when looking at the INT transformed relative abundances as phenotypes (Figure 4C) the peak on chromosome 10 is also present for the abundance of the family *Arcobacteraceae* although no suggestive SNP is present (top SNP, AX-88038371 at position 15,555,246bp). No other suggestive SNPS were present.

Using a list of differentially expressed genes in gill and head-kidney found between AGD susceptible and more resistant fish from a previous study (Robledo et al., 2020) we attempted to identify associated candidate genes for AGD resistance. Looking in a 2Mb range of the suggestive SNPs and the peak on chromosome 10 where the peak is present we identified several genes (Table S2).

Firstly the gene *cullin-4b*, a part of the cullin family of proteins, essential for protein degradation through ubiquitination (Fan et al., 2022; Sarikas et al., 2011) was found on chromosome 4 close to the SNP associated with mean gill score and reported to be more expressed in head-kidney of susceptible fish (Robledo et al., 2020). Secondly the gene *slc4a2* encodes an anion exchange protein involved in exchange of CL- and HCO3-across cell membranes and likely contributes to Ph regulation in fish (Esbaugh et al., 2012; Romero et al., 2004; Shmukler et al., 2008) is located in proximity to the SNP on chromosome 4 associated with alpha diversity (q=0) and reported to be more expressed in resistant fish gills (Robledo et al., 2020). Thirdly, *slc34a2* encodes a phosphate carrier protein involved in cellular homeostasis of inorganic phosphate (Hilfiker et al., 1998; Werner et al., 2016) and was found in proximity to the peak on chromosome 10 and was reported to have higher expression in head-kidney of AGD susceptible fish (Robledo et al., 2020). Finally, *eps15* coding for epidermal growth factor substrate 15, is involved in the internalisation of the epidermal growth factor receptor (EGFR) and secretion and endocytosis in general (Henegouwen & Paul, 2009), was found close to the peak on chromosome 10 and was reported to be more expressed in gill of susceptible fish. Literature from murine models and humans suggests that EGFR mucin is involved in processes surrounding key indicators of AGD including increased mucus production and hyperplasia (Burgel & Nadel, 2004; Takeyama et al., 2008).

## DISCUSSION

In this study we demonstrated the value of applying a holo-omic approach to study AGD resistance in Atlantic salmon. We described the composition of the gill microbiota of AGD affected salmon and characterised the differences in microbiota composition based on the severity of the disease. We then conducted a GWAS with both traditional resistance indicator traits and traits of the gill microbiota composition. Here we discuss the perspectives of the results and address limitations of the study.

### Influence of environmental factors, sampling procedure and *Simkaniaceae* presence

The Atlantic salmon gill microbiota has been suggested to be altered in response to AGD (Birlanga et al., 2022; Clinton et al., 2024; Slinger et al., 2020). While our observations are in agreement with microbiota differences in relation to AGD, the results differ somewhat from the microbiota composition described in other AGD related gill microbiota studies. This difference may be due to more transient environmental factors such as differences in water pH or salinity (Lokesh & Kiron, 2016; Sylvain et al., 2016). However, the differences may also be due to the procedure used to sample the gill microbiota.

In this study we used gill tissue rather than relying on alternative strategies such as gill mucus scrapes or mucosal swabs. These strategies have been reported to be reliable alternatives for assessing gill microbiota in salmonids, but are also reported to have compositional differences when compared to each other (Birlanga et al., 2022; Clinton et al., 2021; Clokie et al., 2022). Although gill tissue is undoubtedly an invasive sampling technique it may be appropriate in certain cases, especially in the case of endosymbiotic bacteria which may reside and propagate in the gill tissue or derived directly from parasites and that would be missed from only sampling a mucosal swap. Perhaps including a combination of mucosal swabs along with whole gill tissue will give a better representation of the entirety of the gill microbiota.

The most striking difference to other studies was the high relative abundance of *Simkaniaceae*. *Candidatus* Syngnamydia Salmonis, a member of the *Simkaniaceae* has been found in association with *Neoparamoeba perurans*, the AGD causing parasite and have even been grown in co-culture with the parasite (A. Nylund et al., 2018; S. Nylund et al., 2015). Moreover *Simkaniaceae* have been reported in other AGD related studies (Birlanga et al., 2022; Clinton et al., 2024), although in lower abundances. Although the representative ASV of the *Simkaniaceae* in this study shows a high percentage identity with *Candidatus* Syngnamydia Salmonis we cannot conclusively confirm its identity. These observations support the previous notion that our results may represent both the internal and external part of the Atlantic salmon gill microbiota along with a potential endosymbiont of the amoebic parasite itself. As suggested in (A. Nylund et al., 2018), *Simkaniaceae* is likely not universal to all *N. perurans* and may thus be opportunists in this regard. The observed correlation between *Simkaniaceae* and amoebic load in this study suggests a potential relationship. However, whether the presence/absence or abundance of this *Simkaniace* has an influence on the virulence of the amoeba or disease progression of AGD – as has been shown with another bacterium (Embar-Gopinath et al., 2005, 2006) – warrants further investigation.

Due to the quantitative study design, a control group of unaffected salmon was not included. The inclusion of a control group would likely give a better clue as to whether *Simkaniaceae* are truly derived from the *P. perurans –* although this is very likely given previous studies. Moreover, a control group would confirm whether the microbiota of AGD affected salmon is more disturbed as compared to an uninfected state.

### Microbiota diversity in relation to AGD scoring methods

The results of the microbiota analysis differed based on the AGD scoring methods. Specifically, one diversity metric differed among samples grouped by amoebic load but not by gill damage and vice versa. This might indicate that the typical resistance phenotypes usually used for AGD may not fully capture the phenotypic nature of the disease and argues for including other layers like that of the microbiota in elucidating resistance mechanisms.

The differences observed among samples grouped by amoebic load in alpha diversity measures indicated that the diversity is reduced in samples with higher amoebic load. This is likely due to certain higher abundance taxa since the differences observed are clearer in the diversity metrics where the abundance of taxa is taken into account. This is supported by the observation that the bacteria belonging to the family *Simkaniaceae* – the most abundant bacteria in our dataset – correlated with amoebic load value (Figure 3B).

### Integrating microbiota traits in GWAS for AGD resistance

A goal of this study was to use a holo-omic approach by conducting a GWAS using both the traditional AGD scoring methods and to include traits of the microbiota composition as resistance phenotypes. It should be noted that the GWAS is underpowered due to the small sample size, which was reflected in the q-q plots of expected vs observed p-values (Figure 4B). Due to unreliable heritability estimates we decided against estimating the percentage of explained variability of each suggestive SNP as the estimates would likely be under/over estimated due to the high SE of heritability and lead to spurious results. Despite a modest sample size and high standard error of heritabilities, our results conform rather nicely with previous studies on AGD resistance as the heritability estimate of the gill score resistance trait was 0.20.

Interpreting microbiota traits in the context of a GWAS on a disease challenged population is complex. While changes in the microbiota between healthy Atlantic salmon gills and AGD infected Atlantic salmon gills are documented (Birlanga et al., 2022; Botwright et al., 2021; Clinton et al., 2024), it is unclear what constitutes a desirable or healthy gill microbiota in context of disease resistance. Is it high or low microbial diversity, a specific microbiota composition, or the presence of a few specific microbes? One possibility is that genetically conveyed resistance allows for the microbiota to stay in a more robust state compared to more susceptible genotypes, suggesting that host genetics influence microbiota composition in a way that enhances resistance. Thus, including traits of the microbiota as covariates with the more conventional resistance traits may add more holistic insights into host genetic-microbiota-parasite interactions compared to traditional GWAS approaches only linking host genotype and phenotype.

### Opportunities and challenges in microbiota-based GWAS

When utilising the relative abundance of individual bacterial families—or any other taxonomic rank—in disease resistance-based GWAS analyses, having a hypothesis-driven rationale for testing a particular trait strengthens the context of analysis. Although exploratory data analysis can also provide valuable insights–as this study shows. For instance, in the case of the relative abundance of *Arcobacteraceae*, the second most abundant bacterial family, a suggestive SNP was identified on chromosome ssa10. This raises the question: is this genetic effect associated with AGD resistance, or does it reflect a different heritable bacterial population trait influenced by host genetics? Conversely, for *Simkaniaceae*, while no suggestive SNP was observed, it may be reasonable to hypothesise that its relative abundance could serve as a reasonably robust resistance indicator trait, given its documented presence in co-culture with the AGD causing parasite (A. Nylund et al., 2018). It has been noted that GWAS studies applying 16S microbiota data may be problematic to some extent, due to the hierarchical nature of taxonomic data and relative abundance of microbial taxa (Awany et al., 2018; Bruijning et al., 2023). That is because the relative abundances of taxa can be represented across multiple levels (Phylum, Family, Genus, Species etc.) and thus each host SNP may affect these levels differently. The interrelatedness across taxonomic levels may therefore obfuscate the potential to directly suggest resistance indicators from a taxon point of view. However, assuming that the composition of the microbiota is influenced by host genetic factors we can argue that using microbiota traits as a health indicator in a quantitative genetics framework better captures genetic influence on the microbiota that directly contributes to resistance mechanisms. Rather than relying solely on the traditional phenotypes, including the microbiota in the resistance phenotype, putatively introduces a unique, differentially nuanced, and integrated view of resistance mechanisms.

### Towards functional insights

In a holo-omic context the compositional taxonomy-centric nature of 16S metabarcoding can be a limiting factor. While a 16S metabarcoding approach provides a birds-eye view over microbiota dynamics, the approach fails to capture information regarding the gene content, functions and activity of the microbiota. Although methods have been developed to infer functional insights based on 16S metabarcoding data (Aßhauer et al., 2015; Douglas et al., 2020; Langille et al., 2013), these methods are constrained in their ability to accurately infer functional capacity (Sevigny et al., 2019), particularly in animal and environmental studies (Sun et al., 2020) or when considering the taxonomic resolution of the 16S amplicon data (Alberdi et al., 2022; Antony-Babu et al., 2017; Welch et al., 2002). The limitations of marker-gene based prediction methods are overcome by Integrating long-read metagenomic sequencing and metatranscriptomics layers. These layers include whole bacterial genomes, associated functional pathways, microbial genetic variants, metabolism predictions and gene expression, all of which may be applied as phenotypes in a GWAS (Doolittle & Booth, 2017; Sanna et al., 2022). Thus, Including these -omic layers, although costly, may give a more clear picture of the functional dynamics involved in coordinating host-microbiota responses such as the gill microbiota of AGD affected salmon and the mechanisms underlying AGD resistance/susceptibility.

Highlighting the added value of including layers from both the host and microbiota, we used a set of differentially expressed genes from the same population (Robledo et al., 2020), based on their location in proximity of the suggestive SNPs. This host-transcriptomic layer can assist in elucidating mechanisms of resistance and identify genes of interest in a putative QTL identified using the layer of the microbiota. This revealed interesting candidate genes involved in cellular processes and homeostasis which have not been documented before in relation to AGD (Boison et al., 2019; Robledo et al., 2020). However, given the limitations of this study regarding sample size and inability to compare differences in gene expression with regard to SNP allelic frequencies, these results should be taken with caution.

### Impact and prospects for future research

The limited knowledge surrounding gill microbiota dynamics and its role in disease as well as the complex nature of AGD and its resistance mechanisms, highlights the need for novel approaches. The idea of incorporating the microbiota to study AGD resistance mechanisms through a holo-omic lens simultaneously poses interesting challenges and questions, which our findings begin to address. The results of the study indicate differences in the microbiota traits based on traditional AGD scoring methods, showcasing the potential of using traits of the microbiota to study AGD disease resistance mechanisms. The techniques applied here can be of value in further understanding AGD resistance or susceptibility. However further larger-scale studies are needed to corroborate the results and draw more confirmed conclusions about the underlying mechanisms governing concerted host-microbiota disease responses.

## DATA AVAILABILITY

Raw 16S sequences for all samples and controls have been deposited at the European Nucleotide Archive, ENA, under reference number PRJEB83575. The scripts and data needed to reproduce the results are available at https://github.com/eirikurandri/AGD_HOL.

## ETHICS STATEMENT

All animals were reared in accordance with relevant national and EU legislation concerning health and welfare. The challenge experiment was performed by the Marine Environmental Research Laboratory (Machrihanish, UK) under approval of the ethics review committee of the University of Stirling (Stirling, UK) and according to Home Office license requirements. Landcatch are accredited participants in the RSPCA Freedom Foods standard, the Scottish Salmon Producers Organization Code of Good Practice, and the EU Code-EFABAR Code of Good Practice for Farm Animal Breeding and Reproduction Organizations.

## AUTHOR CONTRIBUTIONS

**Eiríkur A. Thormar:** Conceptualization; data curation; formal analysis: investigation; methodology; visualization; funding acquisition; writing – original draft; writing – review and editing. **Clémence Fraslin:** data curation; formal analysis; investigation; methodology; software; visualization; writing – original draft; writing – review and editing; supervision. **Morten T. Limborg:** Conceptualization; data curation; funding acquisition; resources; supervision; writing – original draft; writing – review and editing. **Diego Robledo:** Conceptualization; Project administration; supervision; data curation; funding acquisition; resources; writing – review and editing

## ACKNOWLEDGEMENTS

This work was funded by EMBO Scientific Exchange Grant 9514, The Danish National Research Foundation award (CEH – DNRF143). The authors gratefully acknowledge funding from Innovate UK and BBSRC (BB/M028321/1). DR was supported by BBSRC Institute Strategic Grants to the Roslin Institute (BBS/E/20002172, BBS/E/D/30002275, BBS/E/D/10002070 and BBS/E/RL/230002A), and by the Oportunius programme of the Axencia Galega the Innovación (GAIN, Xunta de Galicia).

## Notes

### Competing Interest Statement

The authors have declared no competing interest.

