## Supplementary file S1 for "Amoebic gill disease (AGD) in Atlantic salmon investigated through a holo-omic lens"

**Supplementary figures and tables:**

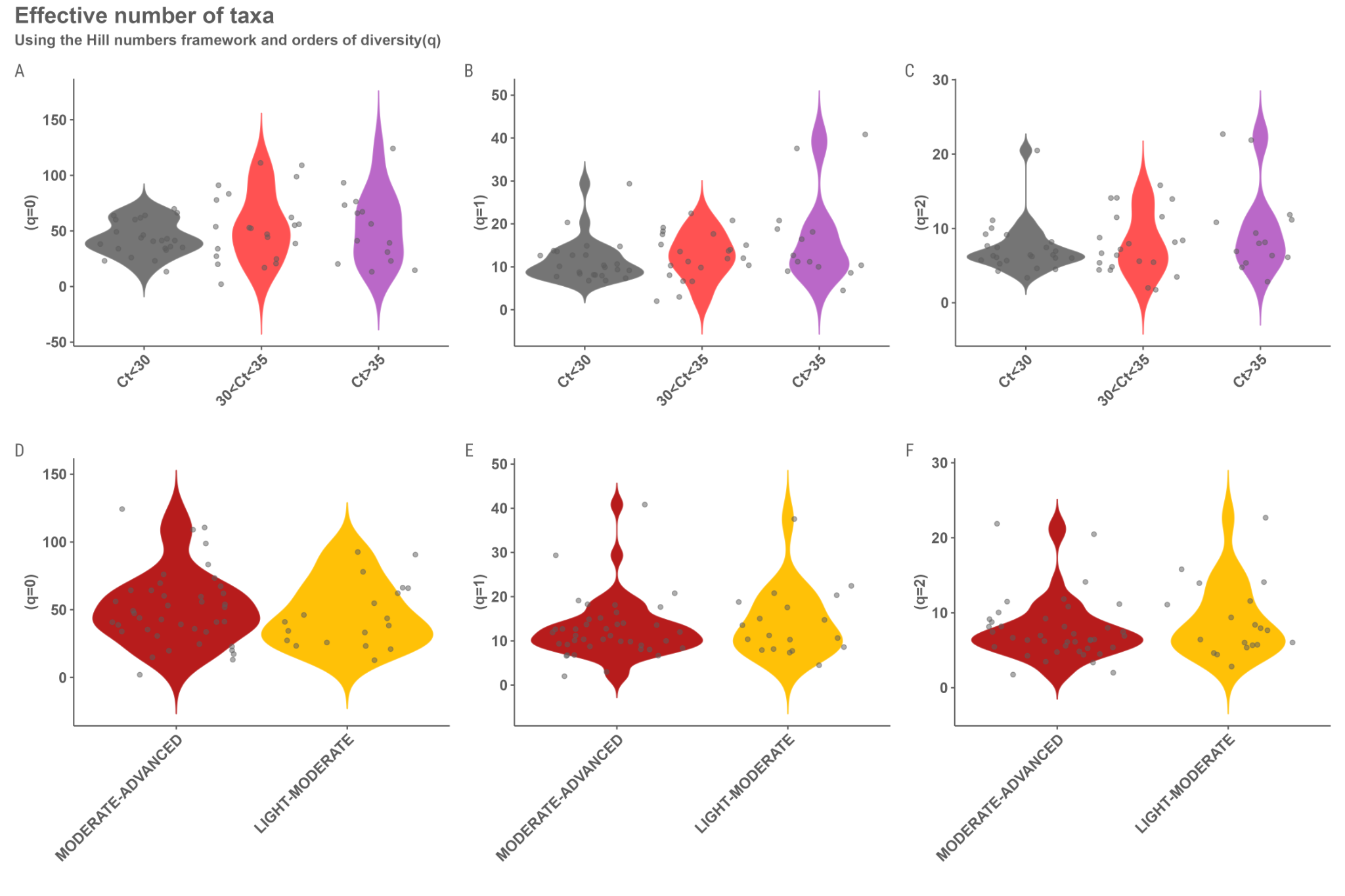

**Figure S1:** ASV level *comparison of alpha diversity estimates based on hill numbers order of diversity q=0, q=1 and q=2 among samples grouped by A-C) amoebic load Ct value where the colours represent Ct<30 (grey), 30<Ct>35 (pink) and Ct>35 (purple), and D-F) gill damage where the colours represent the visually scored gill damage groups, Light-Moderate (yellow) and moderate-advanced (red). No significant differences were observed by grouping on this level.*

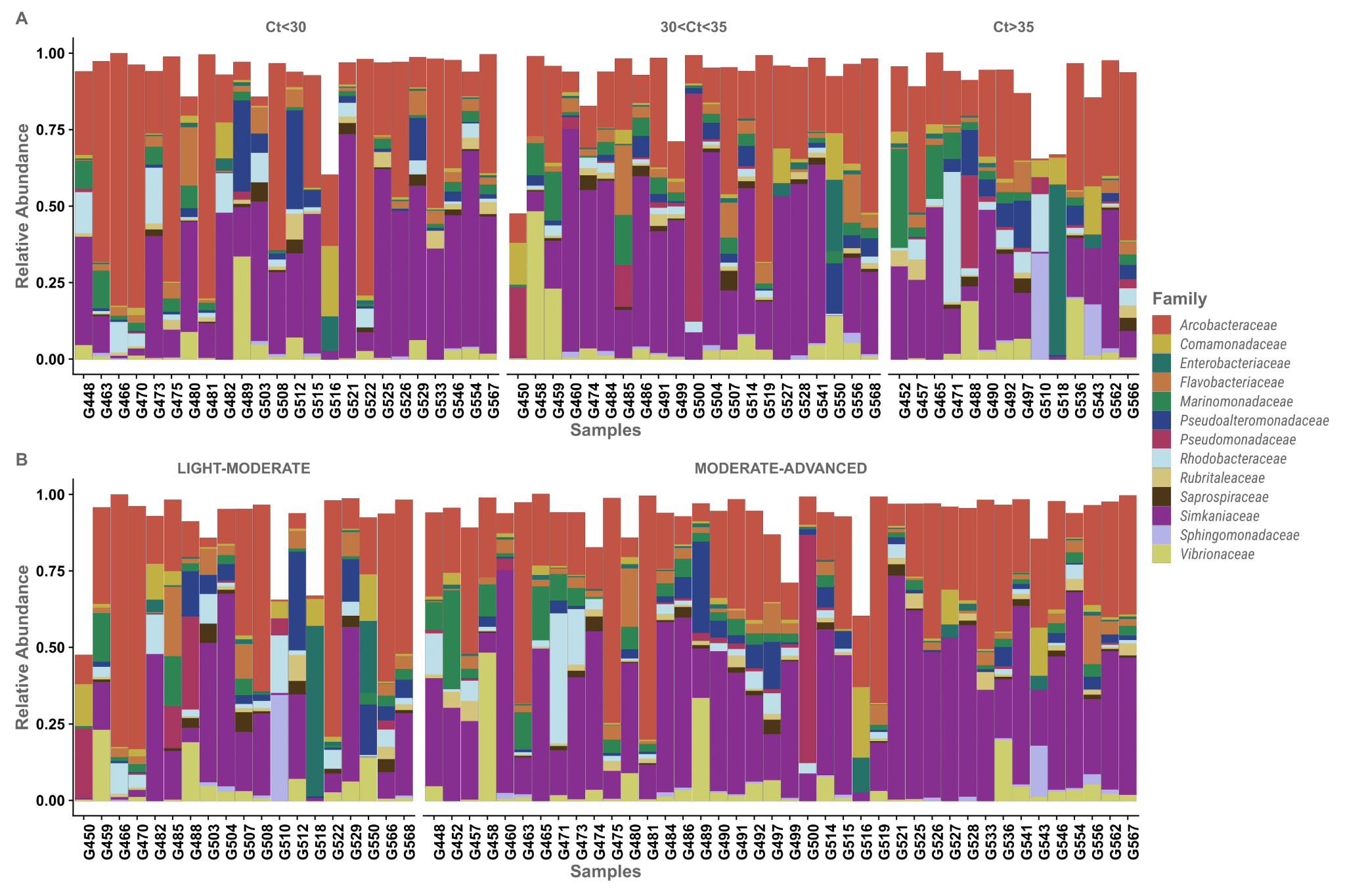

**Figure S2:** *Relative abundances of Bacterial families constituting over 1% of the overall abundance among A) Samples grouped by amoebic load (Ct value) and B) Gill damage. The colors represent the 13 different bacterial families.*

| **Figure S3:** *GWAS for AGD resistance using all phenotypes. A) Amoebic load, B) Gill score, C) Beta diversity axis.1, D) Beta diversity axis.2, E) Alpha diversity (q=0), F) Alpha diversity (q=1), G) Alpha diversity (q=2) and INT transformed relative abundance of bacterial families H) Simkaniaceae, I) Arcobacteracea, J) Vibrionaceae, K) Marinomonadaceae, L) Rhodobacteraceae, M) Pseduoalteromonadaceae and N) Flavobacteraceae. The horizontal red line represents the genome wide significance threshold and the blue line represent the suggestive threshold. The large pink points represents suggestive SNPs.* | |
| --- | --- |
| 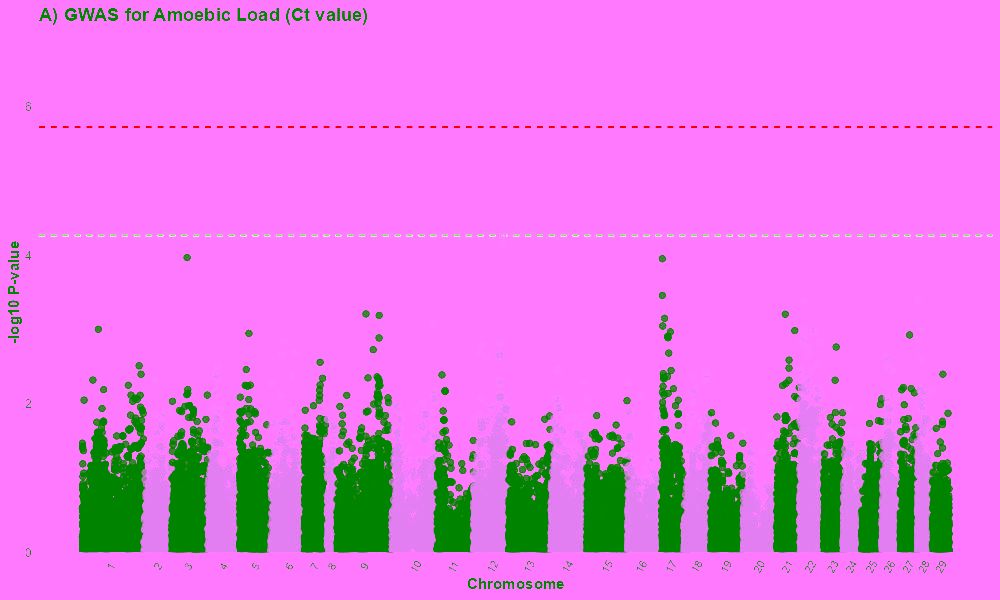 | 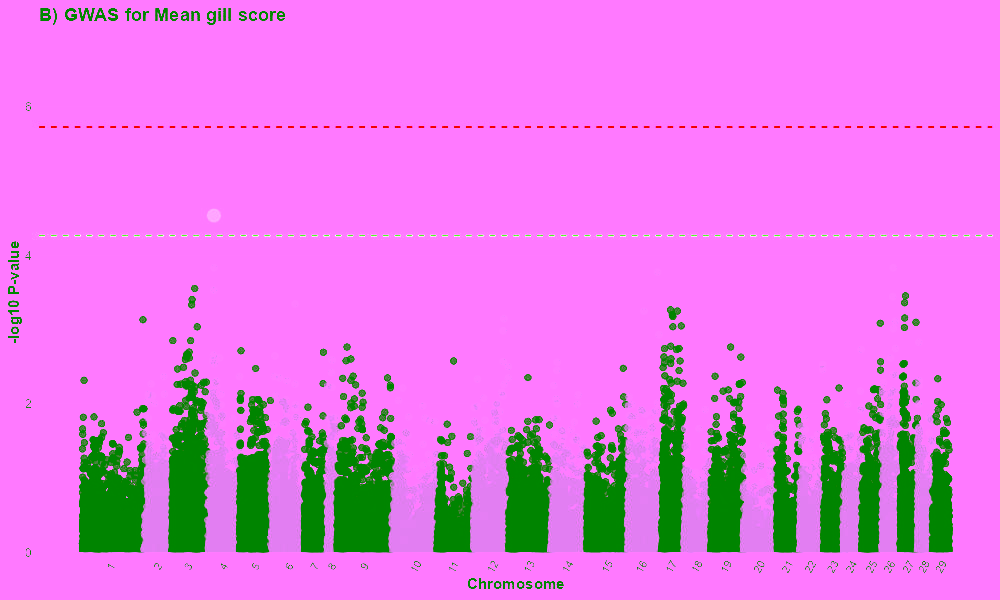 |
| 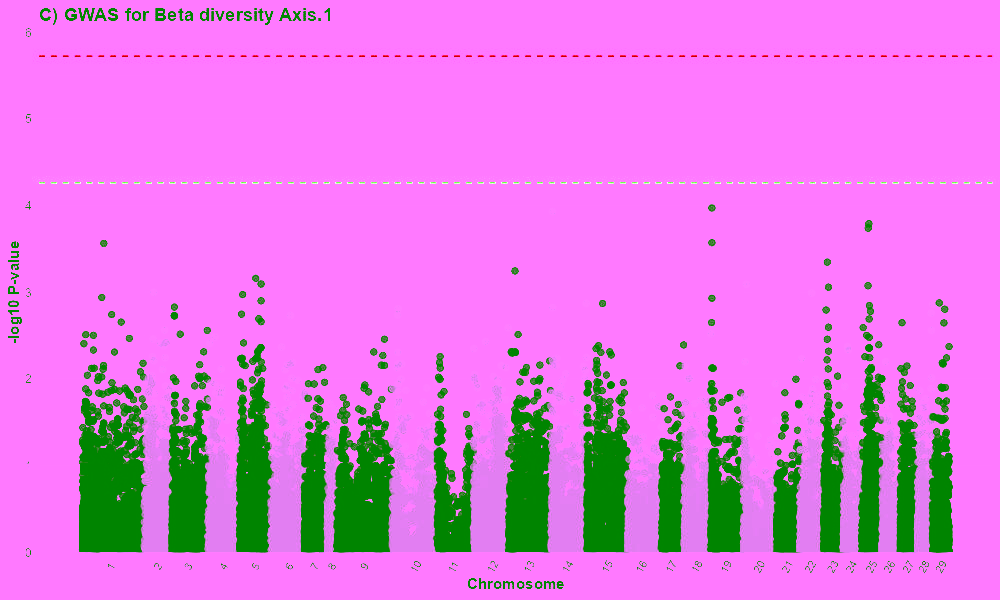 | 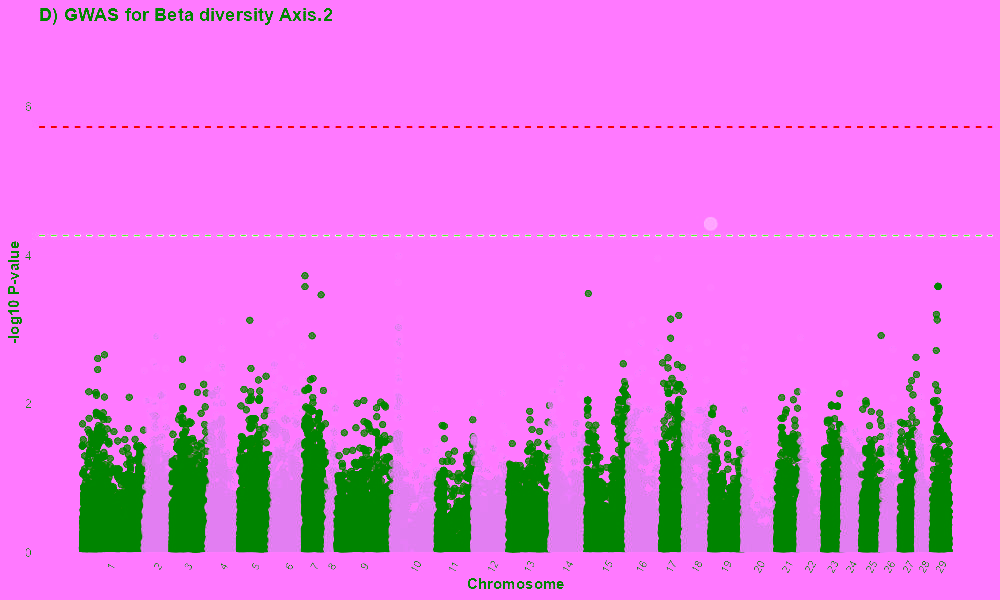 |
| 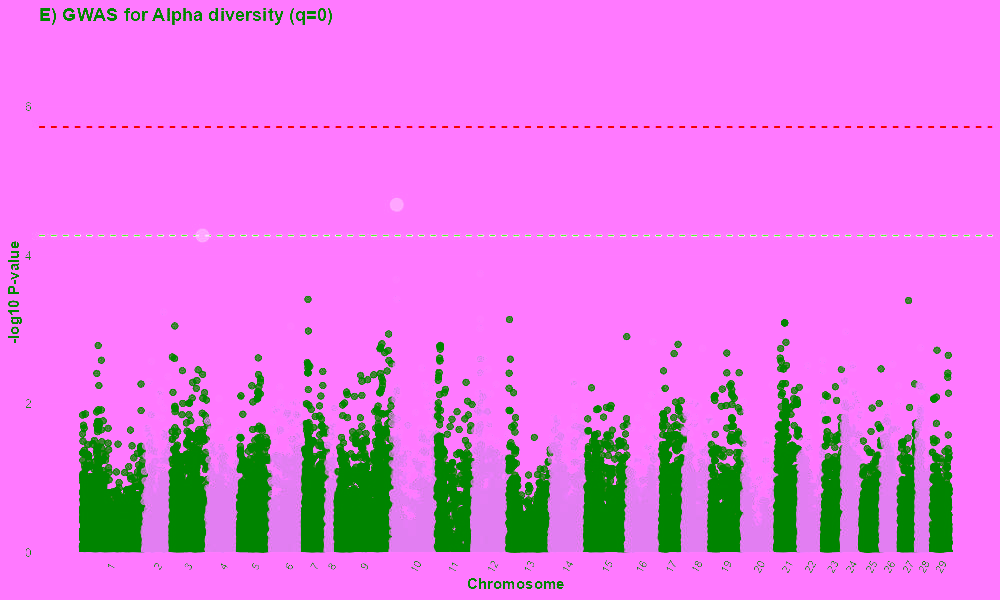 | 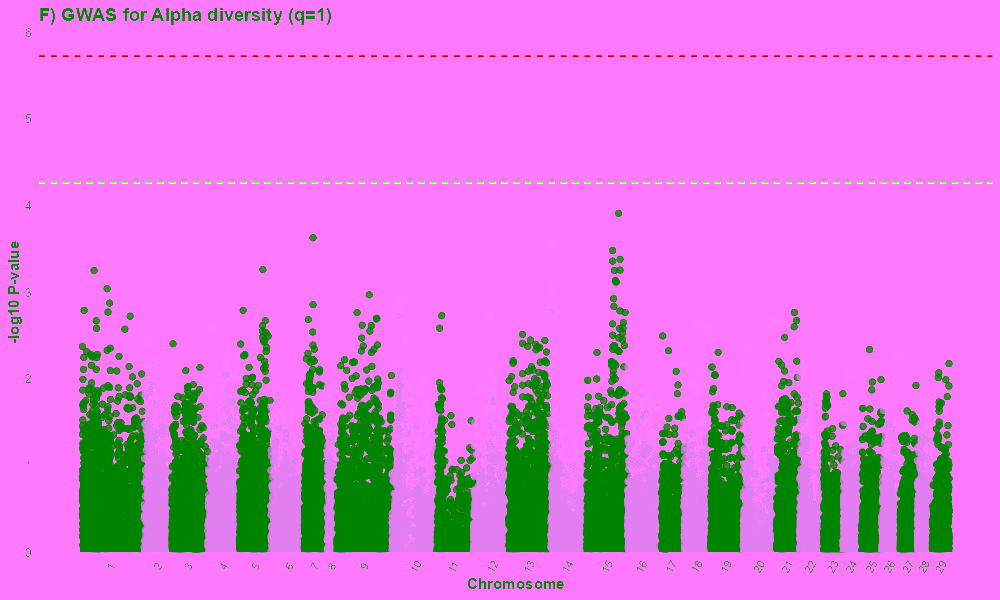 |
| 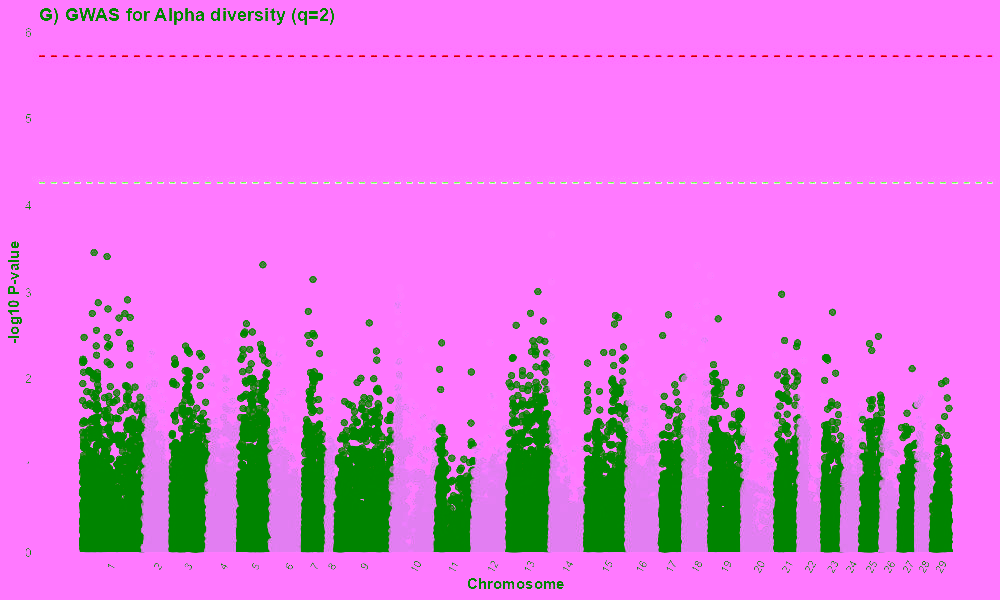 | 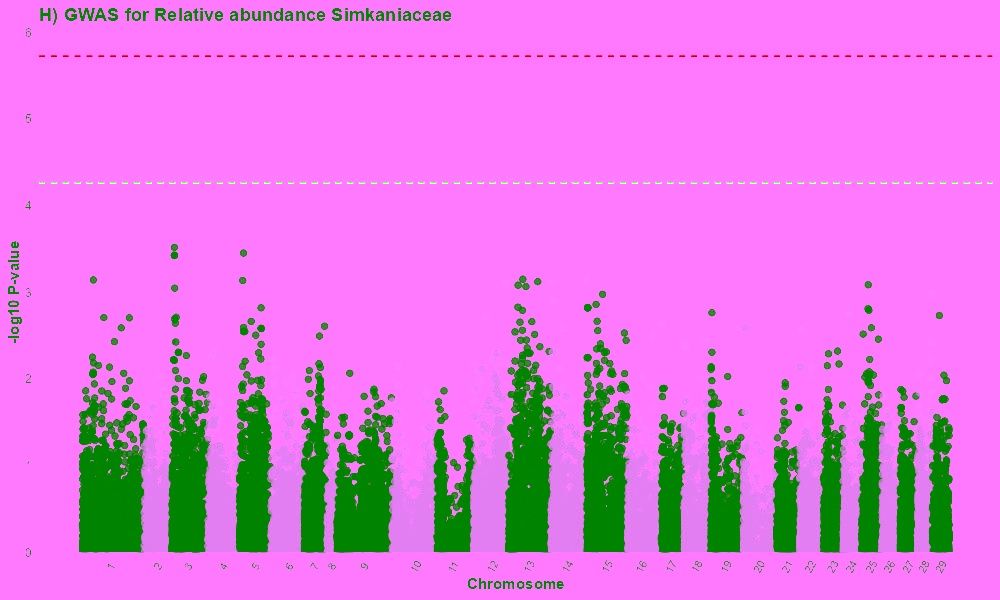 |
| 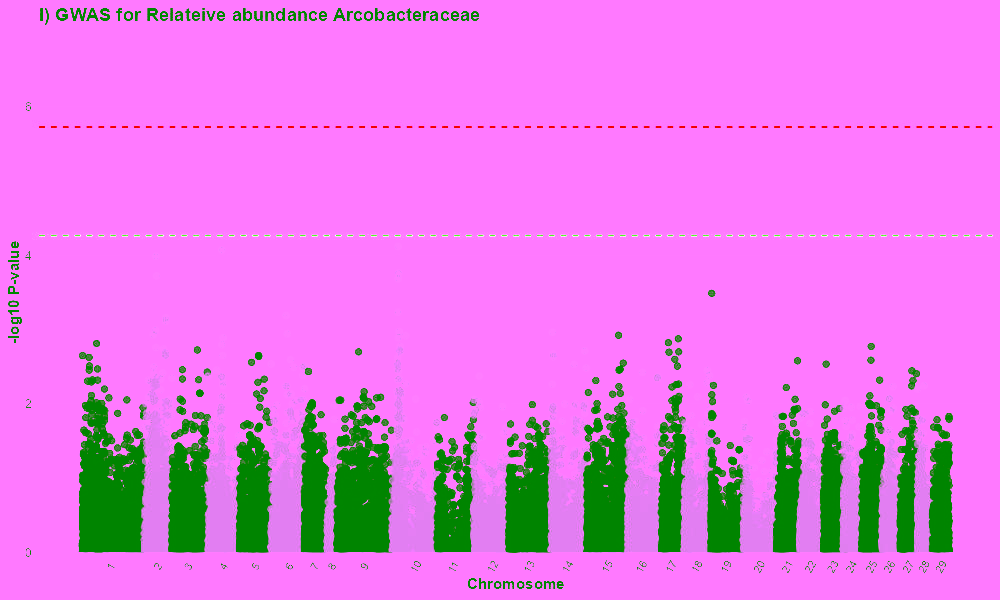 | 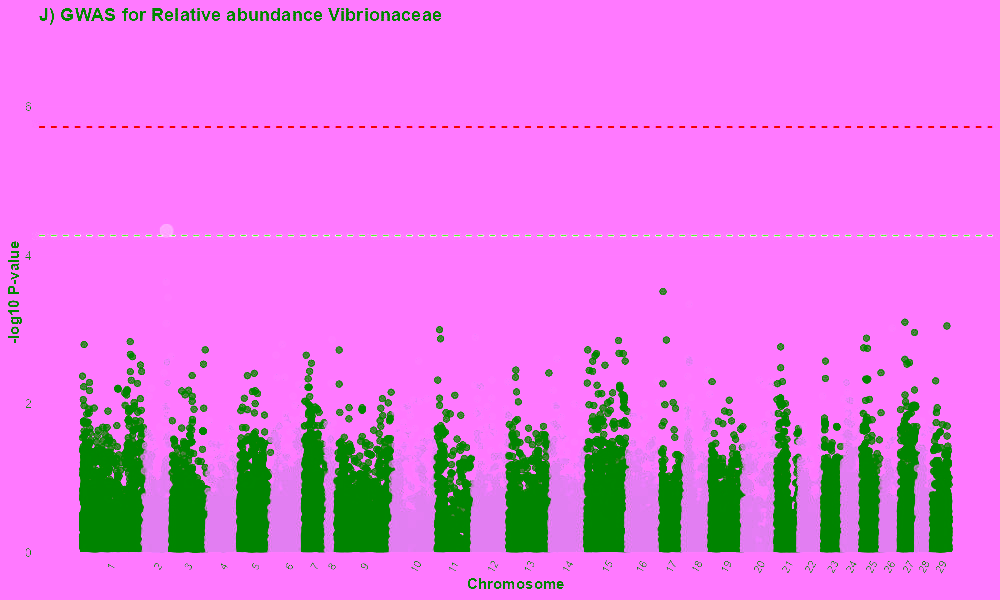 |
| 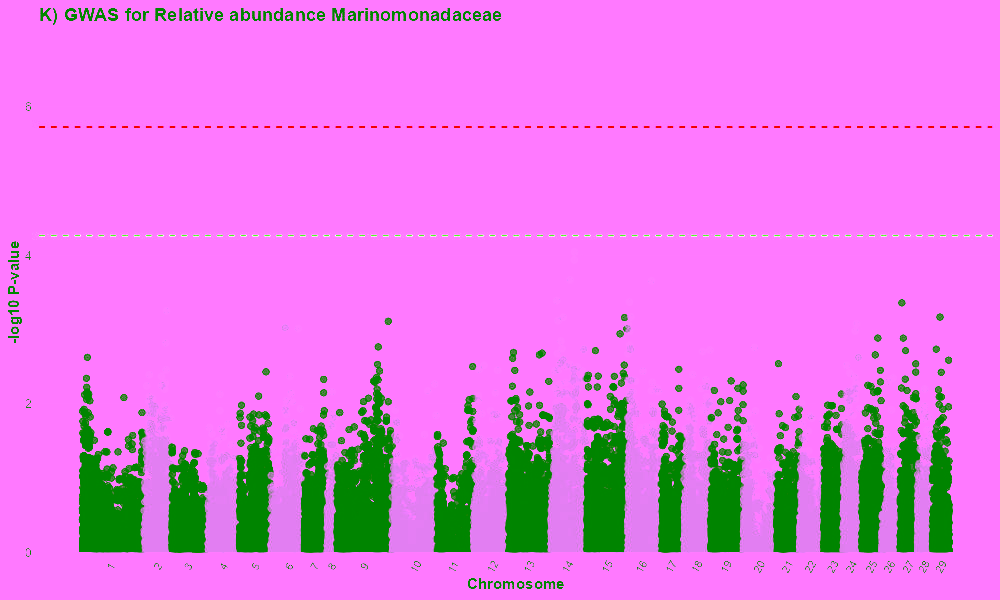 | 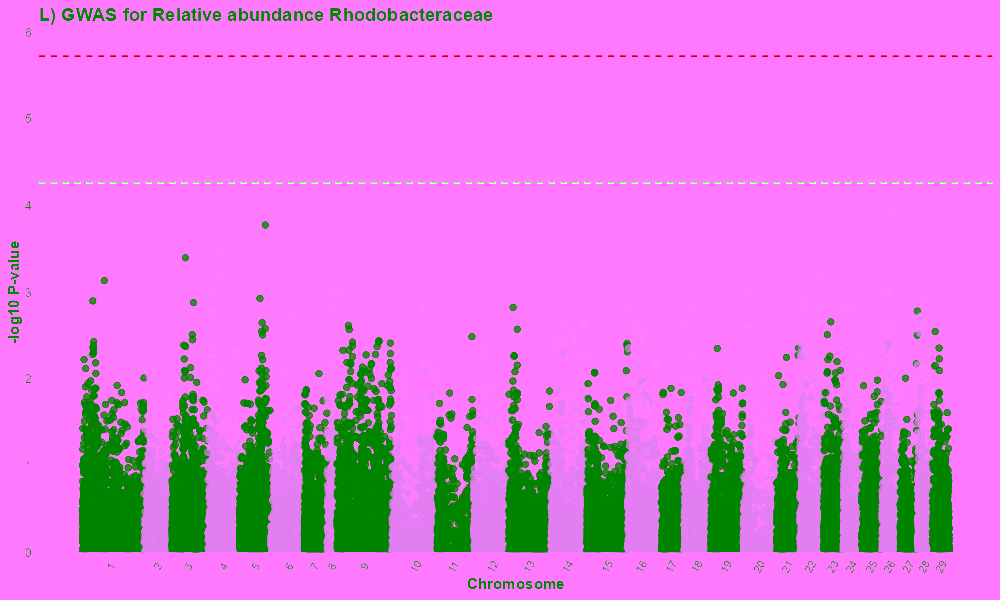 |
| 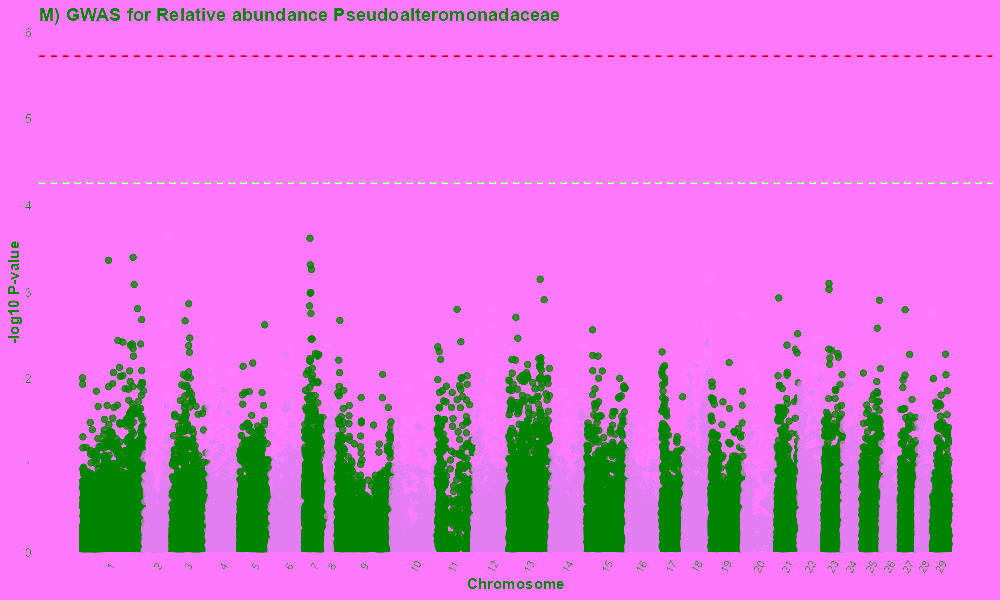 | 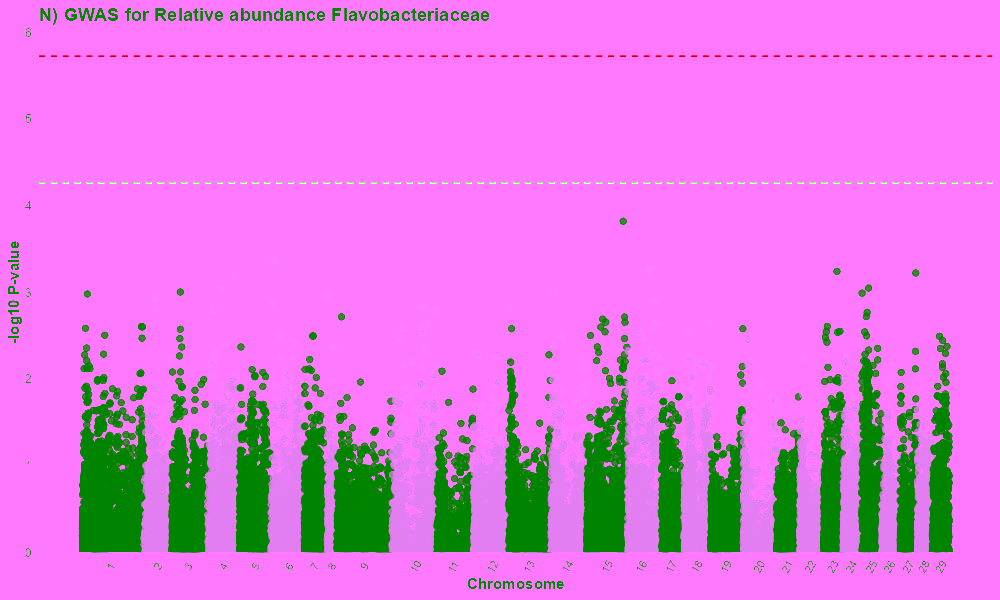 |

| **Table S1:** Variance component estimates for AGD resistance for phenotypes tested and used in subsequent GWAS. | | | | |
| --- | --- | --- | --- | --- |
| **Phenotype** | $\sigma_{a}^{2}(\pm se)$ | $\sigma_{e}^{2}(\pm se)$ | $\sigma_{p}^{2}(\pm se)$ | $h^{2}(\pm se)$ |
| **Gill score** | *0.07 ± 0.1* | *0.26 ± 0.11* | *0.32 ± 0.06* | *0.2 ± 0.32* |
| **Beta diversity axis.2** | *0.01 ± 0.01* | *0.03 ± 0.01* | *0.04 ± 0.01* | *0.17 ± 0.35* |
| **Alpha diversity(q=0)** | *13.21 ± 31.44* | *77.78 ± 33.09* | *90.99 ± 17.12* | *0.15 ± 0.34* |
| **Alpha diversity(q=1)** | *2.08 ± 1.59* | *1.83 ± 1.34* | *3.91 ± 0.77* | *0.53 ± 0.36* |
| **Alpha diversity(q=2)** | *0.84 ± 0.78* | *1.19 ± 0.7* | *2.03 ± 0.39* | *0.41 ± 0.36* |
| **Relative abundance Arcobacteraceae** | *0.19 ± 0.3* | *0.82 ± 0.32* | *1.01 ± 0.19* | *0.18 ± 0.29* |
| **Relative abundance *Marinomonadaceae*** | *0.37 ± 0.37* | *0.5 ± 0.33* | *0.87 ± 0.17* | *0.42 ± 0.39* |
| **Relative abundance *Rhodobacteraceae*** | *0.08 ± 0.23* | *0.82 ± 0.27* | *0.89 ± 0.17* | *0.08 ± 0.26* |
| **Relative abundance *Pseduoalteromonadaceae*** | *0.06 ± 0.27* | *0.83 ± 0.3* | *0.89 ± 0.17* | *0.07 ± 0.3* |
| **Relative abundance *Flavobacteraceae*** | *0.09 ± 0.28* | *0.85 ± 0.31* | *0.94 ± 0.18* | *0.1 ± 0.29* |
| **Relative abundance Arcobacteraceae*** | *0.01 ± 0.02* | *0.03 ± 0.02* | *0.05 ± 0.01* | *0.31 ± 0.32* |
| $\boldsymbol{\sigma}_{\boldsymbol{a}}^{\boldsymbol{2}}\boldsymbol{(\pm}\boldsymbol{se}\boldsymbol{)}$*:The genetic variance,*$\boldsymbol{\sigma}_{\boldsymbol{p}}^{\boldsymbol{2}}\boldsymbol{(\pm}\boldsymbol{se}\boldsymbol{)}$*:The phenotypic variance,*$\boldsymbol{\sigma}_{\boldsymbol{e}}^{\boldsymbol{2}}\boldsymbol{(\pm}\boldsymbol{se}\boldsymbol{)}$ *the residual variance,*$\boldsymbol{h}^{\boldsymbol{2}}\boldsymbol{(\pm}\boldsymbol{se}\boldsymbol{)}$*:the heritability estimate*  **The normalised but non INT transformed relative abundance. Due to the skewness of the data we focused on the INT transformed relative abundance in the study* | | | | |

| **Table S2:** Differentially expressed genes from fish in the same population, form (Robledo et al. 2020), found in 2MB range of the suggestive SNPs and the peak on chromosome 10. | | | | | | |
| --- | --- | --- | --- | --- | --- | --- |
| **Gene symbol** | **Chr.** | **start(bp)** | **end(bp)** | **Annotation** | **Tissue** | **Higher expression** |
| none | 4 | 16,146,246 | 16,147.006 | uncharacterized LOC106582949 | Gill | resistant |
| ***cul4b*** | 4 | 16,981,360 | 17,073,016 | cullin-4B | headkidney | susceptible |
| ***slc4a2*** | 3 | 76,856,731 | 76,860,694 | band 3 anion exchange protein | Gill | resistant |
| ***slc34a2*** | 10 | 10,455,389 | 10,547,023 | phosphate carrier protein | headkidney | susceptible |
| ***eps15*** | 10 | 16,619,912 | 16,655,629 | epidermal growth factor receptor substrate 15 | Gill | susceptible |

| **Table S3:** Taxonomic identity of removed contaminant ASVs | | | |
| --- | --- | --- | --- |
| **Contaminant ASV** | **Class** | **Family** | **Genus** |
| ASV_63 | *Alphaproteobacteria* | *Beijerinckiaceae* | *Methylobacterium-Methylorubrum* |
| ASV_83 | *Alphaproteobacteria* | *Beijerinckiaceae* | *Methylobacterium-Methylorubrum* |
| ASV_94 | *Gammaproteobacteria* | *Moraxellaceae* | *Enhydrobacter* |
| ASV_115 | *Gammaproteobacteria* | *Moraxellaceae* | *Enhydrobacter* |
| ASV_136 | *Gammaproteobacteria* | *Moraxellaceae* | *Acinetobacter* |
| ASV_452 | *Actinobacteria* | *Corynebacteriaceae* | *Corynebacterium* |
| ASV_602 | *Actinobacteria* | *Corynebacteriaceae* | *Corynebacterium* |
